## Supplementary Figures and Results for "A semiconductor 96-microplate platform for electrical-imaging based high-throughput phenotypic screening"

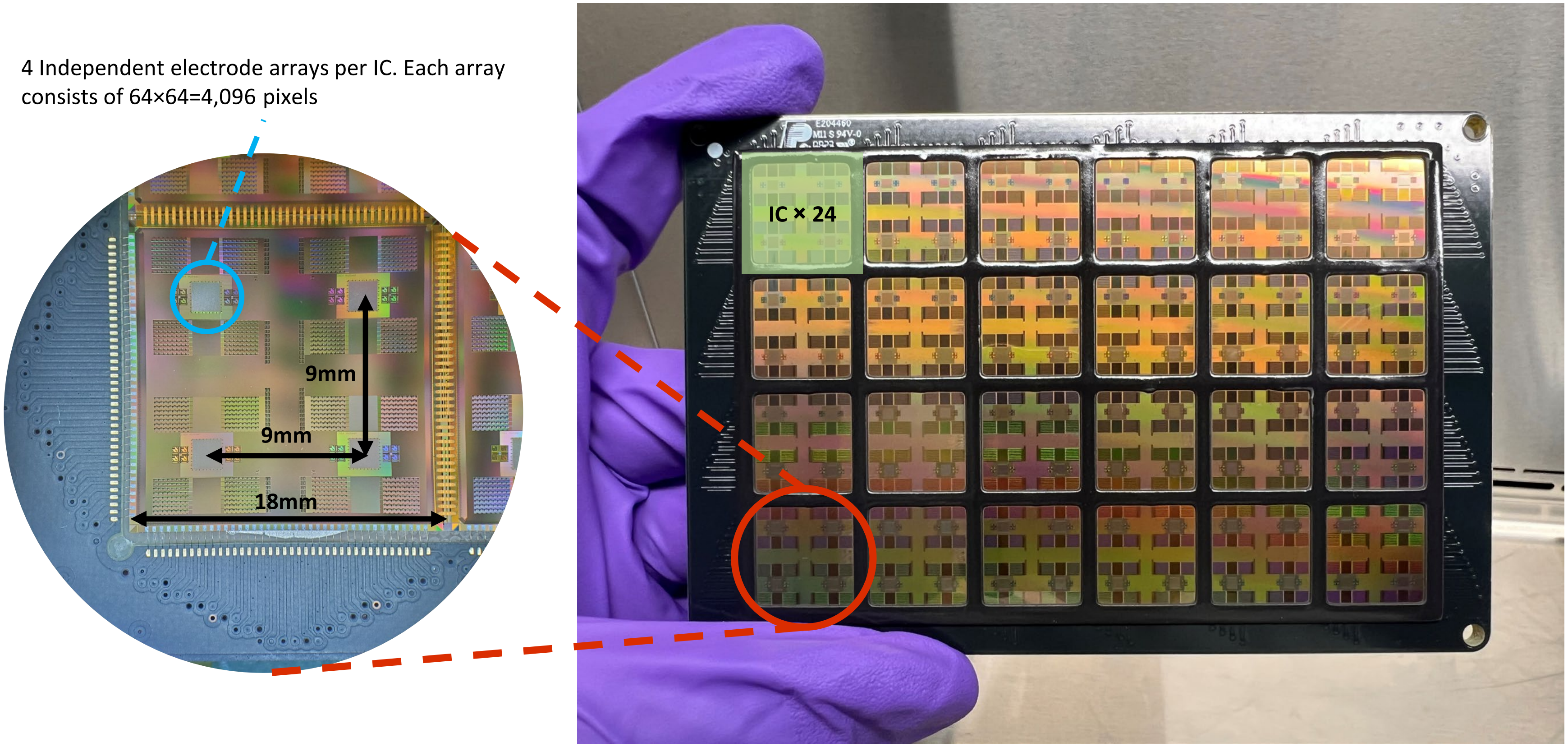


**Supplementary Fig. 1 | Layout and assembly of the semiconductor 96-microplate.** The microplate assembly consists of 24 identical integrated circuits (ICs) interconnected with each other in a 4×6 array, with the peripheral ICs bonded to a PCB for signal route out. The IC is designed to be 18 mm × 18 mm and contains 4 independent electrode arrays at a 9 mm center-to-center spacing. This layout method gives the entire assembly 96 independent electrode arrays with precise array-to-array spacing of exactly 9 mm, the same as a standard 96-microplate. Each IC connects to its neighbor IC via gold wirebonds or redistribution layer (RDL). The chip-to-chip connections allow signals to pass from peripheral ICs to internal ICs, minimizing the number of digital control signals needed for microplate programming.

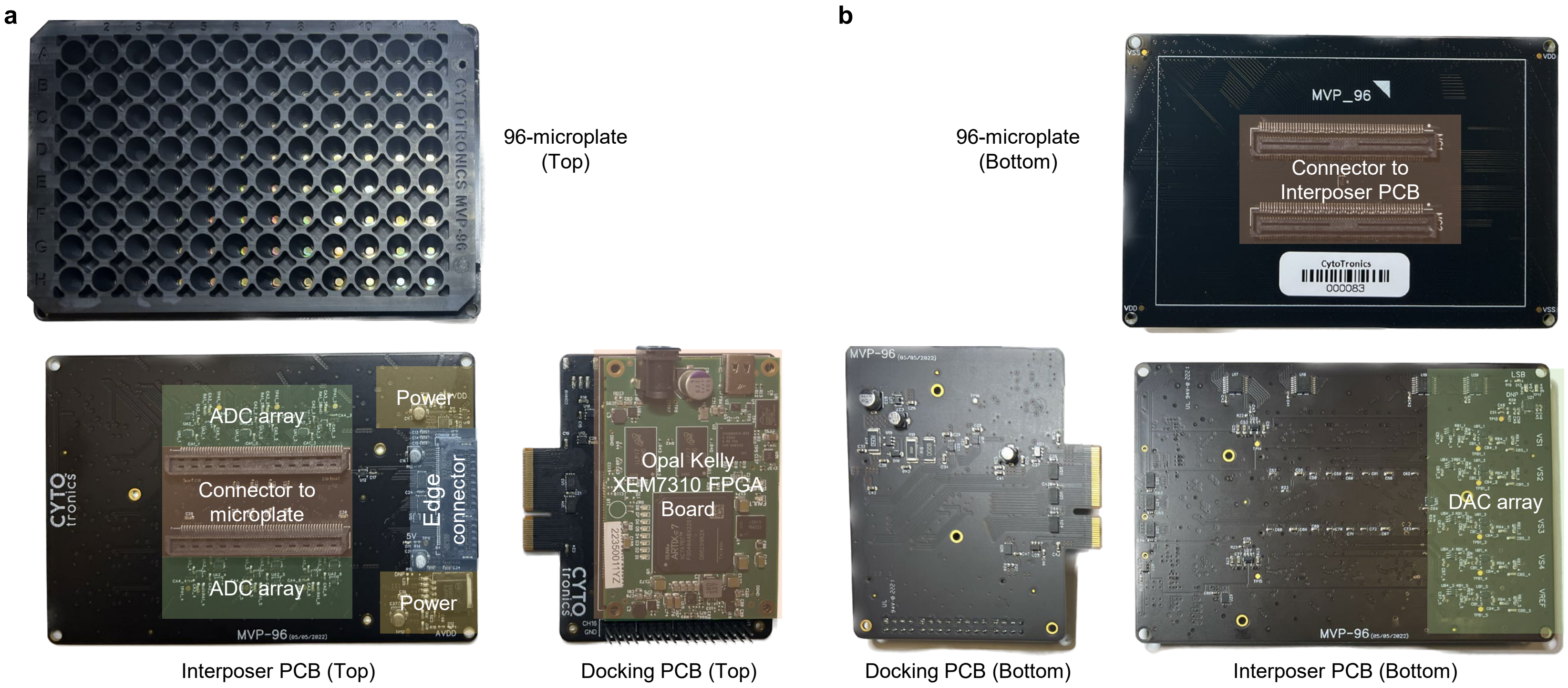


**Supplementary Fig. 2 | Supporting electronics for the semiconductor 96-microplate. a-b,** Top-down (a) and bottom up (b) views of the 96-microplate, Interposer PCB, and Docking PCB. A connector on the backside of the 96-microplate plugs into the Interposer PCB. An edge connector then connects the Interposer PCB and Docking PCBs. An Opal Kelly FPGA board (XEM7310) also plugs into the Docking PCB and creates the universal serial bus (USB) interface to a computer for data acquisition.

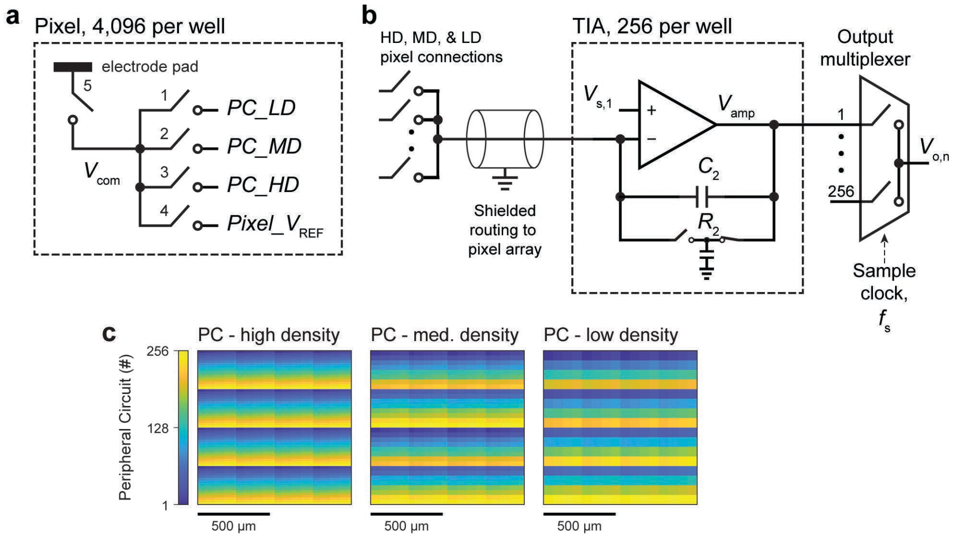


**Supplementary Fig. 3 | Electronic pixel and peripheral circuit schematic for scanned impedance sensing. a,** Five bits of memory per pixel switch the electrode to three options of peripheral circuits (PCs) for measurement or a reference voltage, $Pixel\_V_{ref}$. **b**, Each of the 96 wells contains 256 peripheral circuits configured as transimpedance amplifiers (TIAs) that can connect to a variety of pixels within the pixel array. Each of the 256 peripheral circuits per well are routed to an output pin through a multiplexer. The reference voltages, $V_{s,1}$ and $Pixel\_V_{ref}$, is provided by a digital to analog converter (DAC) and the output, $V_{o,n}$, is sampled by an analog to digital converter (ADC), both located on the Interposer PCB. **c**, High density (HD), medium density (MD), and low density (LD) options are available for scanning across the 64×64 = 4,096 pixels. Digital signals for the scanning are generated by the Opal Kelly XEM7310 FPGA board.

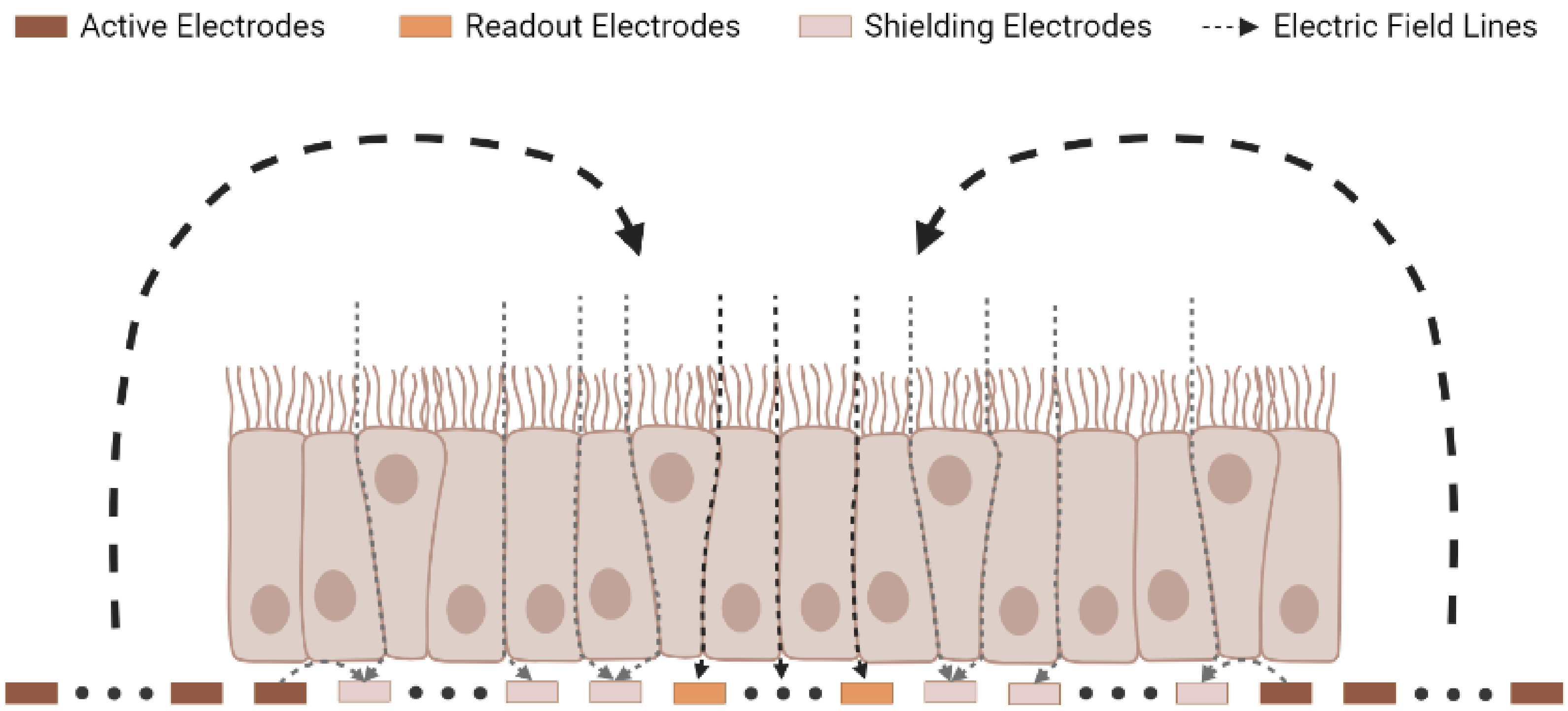


**Supplementary Fig. 4 | Multi-site vertical field biasing scheme allows concurrent measurement from center 4×4=16 electrodes cluster.** A center 4×4=16 electrodes are biased as readout electrodes, with 6 shielding electrodes from each edge of the readout electrodes (total of 240 shielding electrodes) biased the same way as the readout electrodes. The rest of the 3,840 electrodes are active electrodes delivering multi-frequency electric field into the cell culture. The existence of shielding electrodes blocks lateral field component from returning to the readout electrodes and thus they only capture the vertical field component of the electric field. The presence and different activities of cells affect the signal magnitude and phase returning to the readout electrodes. After each frame of measurement, the readout and shielding electrode group shift at a step of 4 electrodes to execute next frame of measurement. A total of 256 frames of measurements constitute one image.

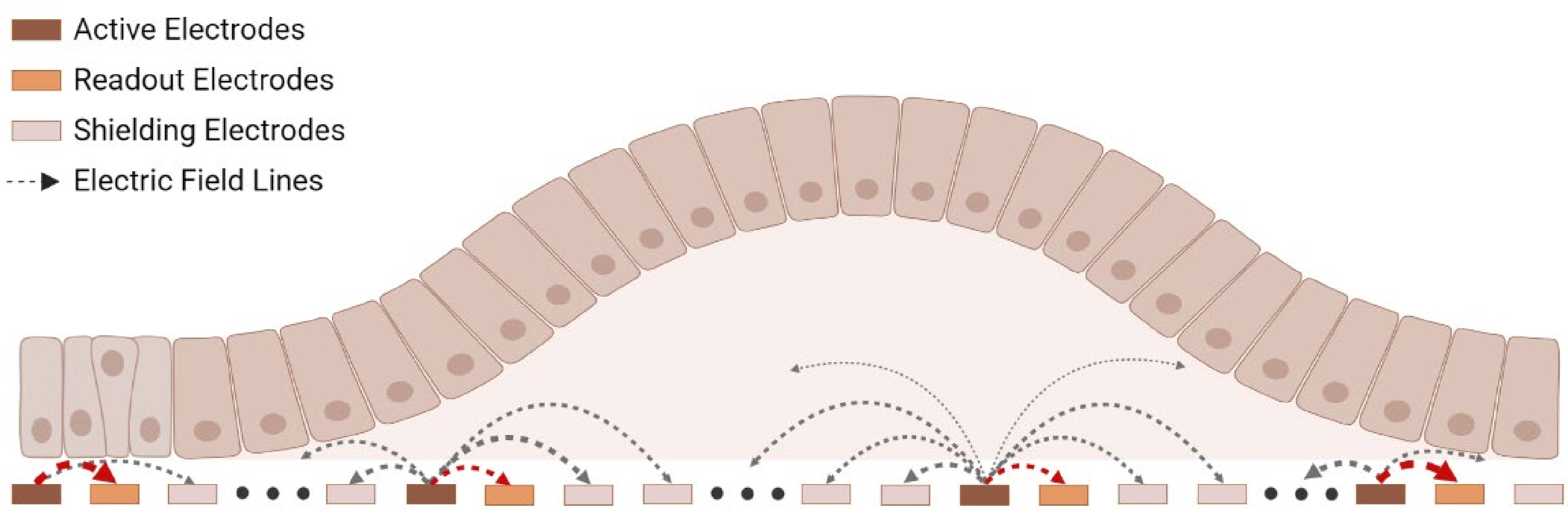


**Supplementary Fig. 5 | Multi-site lateral field biasing scheme allows concurrent measurement from 16 electrodes distant from each other.** 16 out of 4,096 available electrodes are biased as active electrodes where multi-frequency electric field are generated into the cell culture. Each active electrode has 2 return electrodes (total of 32) in orthogonal directions capturing the return signal of the electric field. The presence and different activities of cells disturb the electric field differently and thus affect the return signal magnitude and phase. The 16 active-return electrode groups are separated by at least 12 layers of shielding electrodes to prevent cross-talks interference between electrode groups. After each frame of measurement, the 16 electrode groups shift to a different location (non-repeated locations) to execute next frame of measurement. A total of 256 frames of measurements constitute one image.

**a**

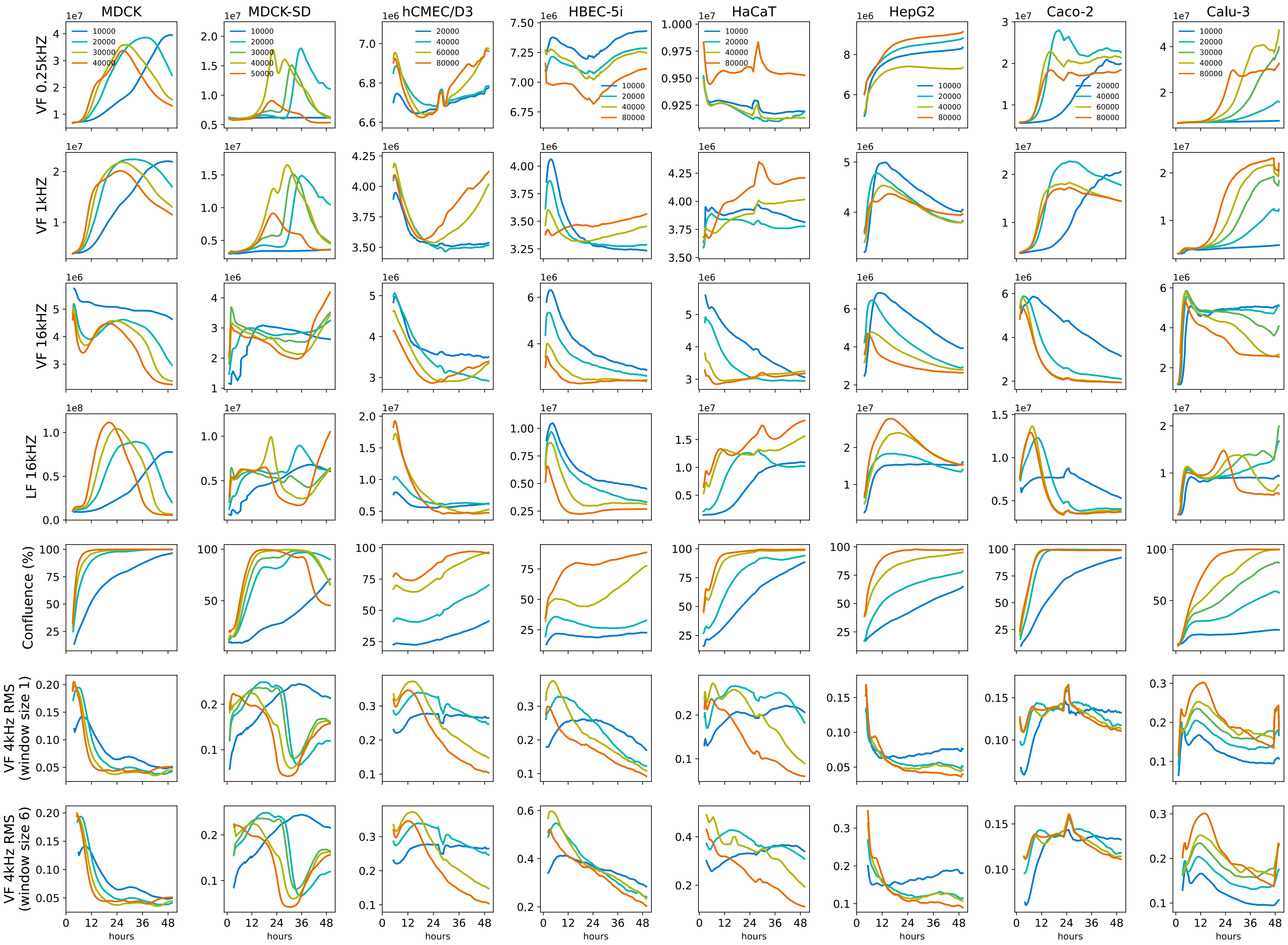


**Supplementary Fig. 6 | Expanded comparison of cell types shown in Fig. 3 and Supplementary Video 1. a,** 8 of the 16 tested cell-types. The total number of cells plated per well for each of cell type is indicated in the legend. Media changes are performed either at 24 or 48 hours depending upon cell type and experiment. The y-axis is scaled on a per cell type basis. Real-time videos of each cell type and plating density are shown in Supplementary Video 1.

b

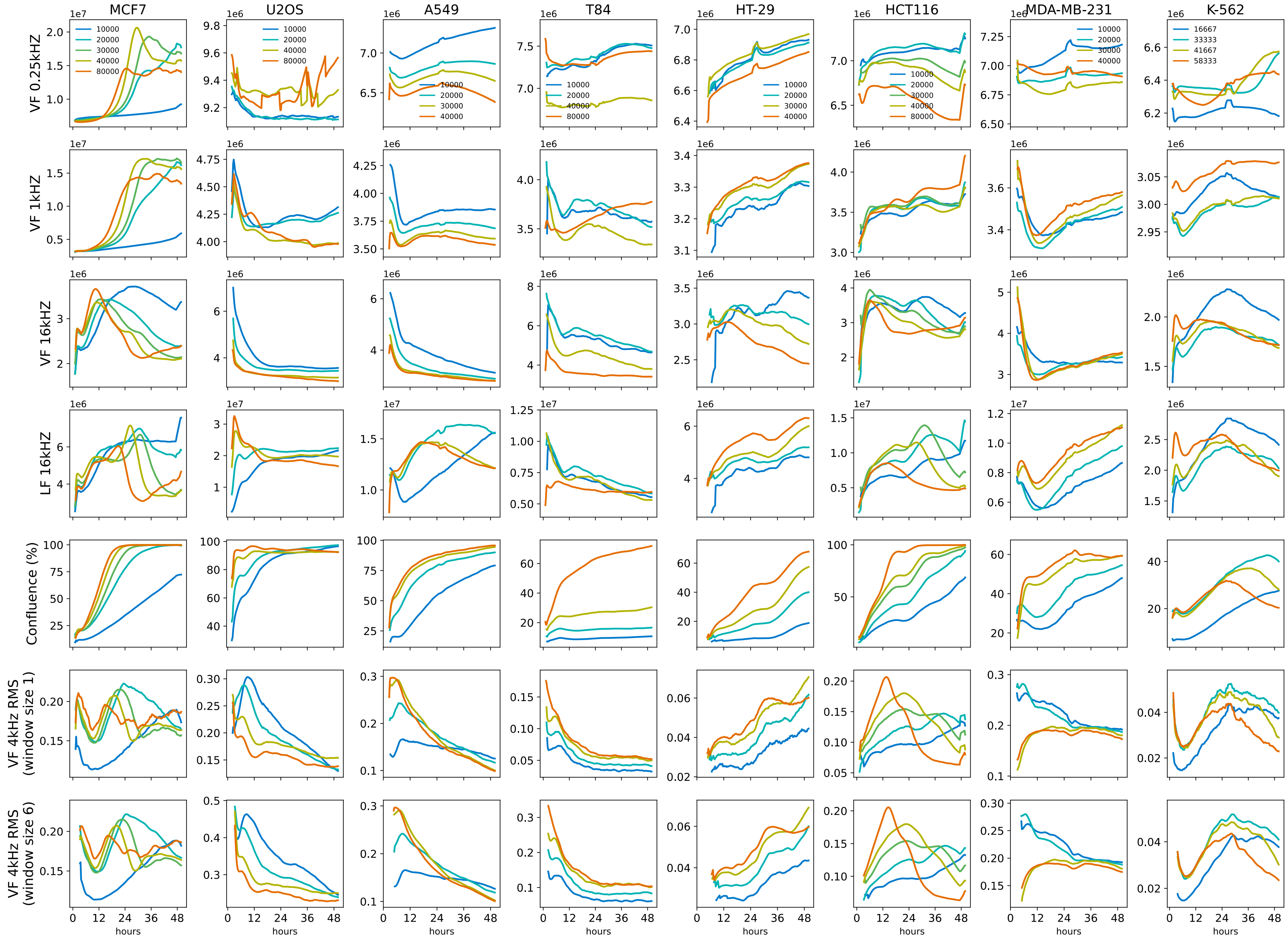


**Supplementary Fig. 6 | Expanded comparison of cell types shown in Fig. 3 and Supplementary Video 1. b,** Remaining 8 tested cell-types. The total number of cells plated per well for each of cell type is indicated in the legend. Media changes are performed either at 24 or 48 hours depending upon cell type and experiment. The y-axis is scaled on a per cell type basis. Real-time videos of each cell type and plating density are shown in Supplementary Video 1.

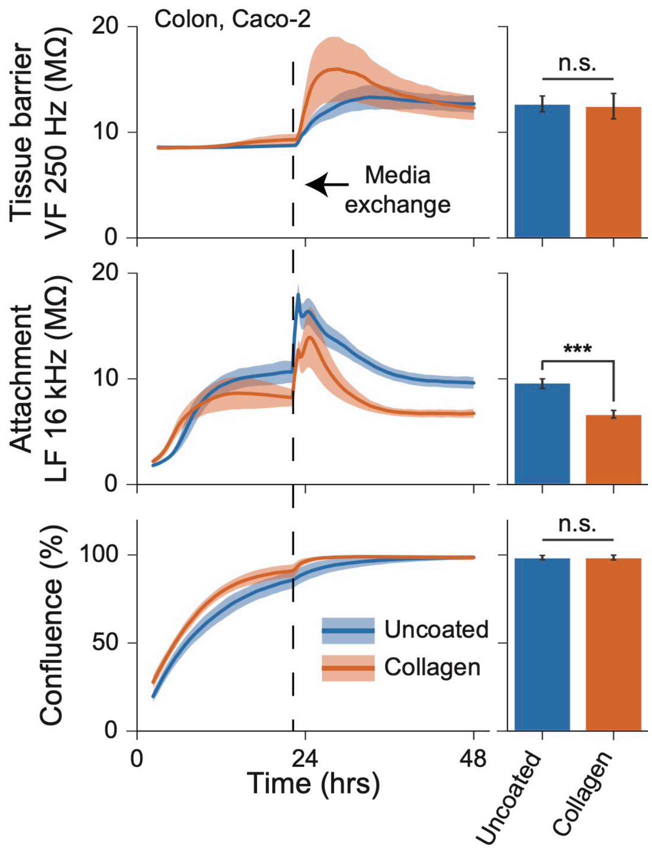


**Supplementary Fig. 7 | Caco-2 cells on Collagen.** A surface coating comparison study shows a difference in Caco-2 cell function from 0 - 48 hours after seeding. Tissues on coated surfaces increased in tissue barrier (VF 250 Hz) more significantly with a media exchange and showed less attachment (LF 16 kHz). Line traces represent mean ± s.d. for 9 wells per condition; bar graphs are at 48 hours (***, $p < 0.0001$, n = 9 wells).

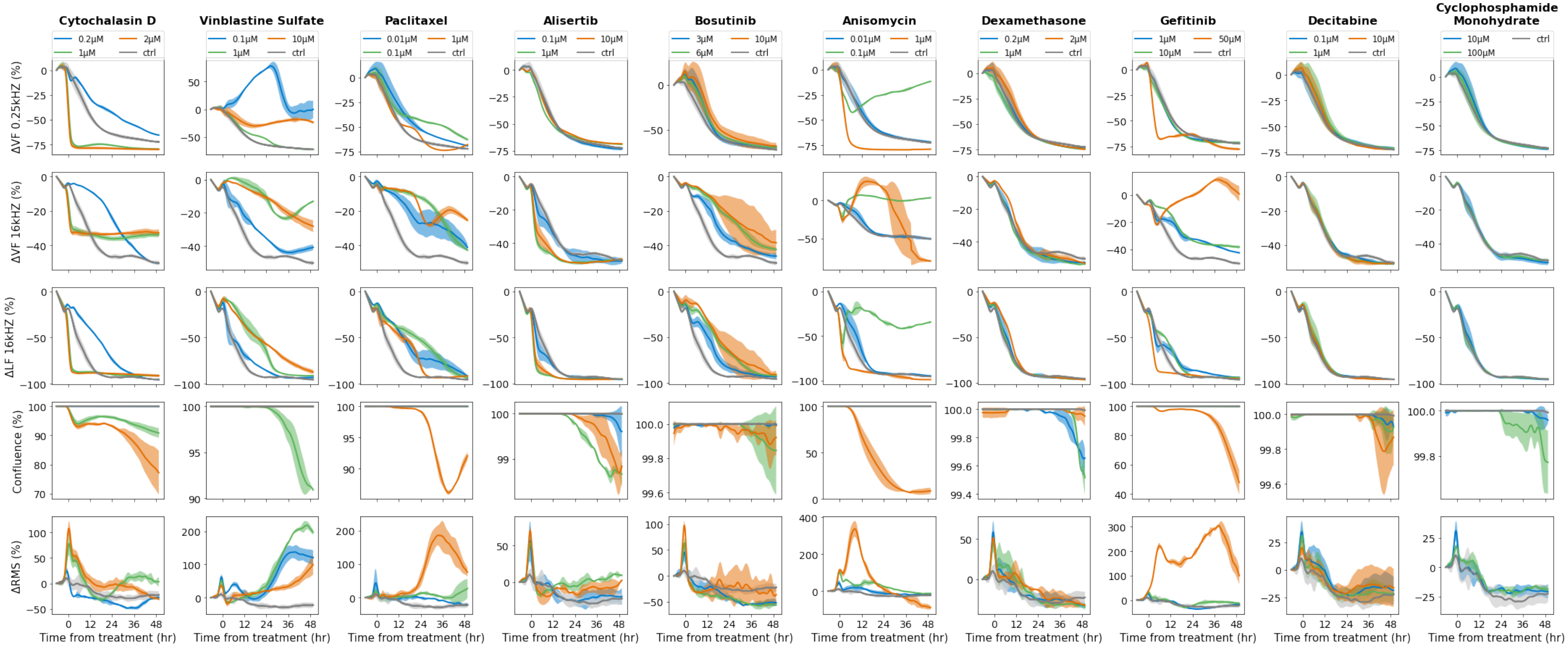


**Supplementary Fig. 8 | Compound applications to MDCK cells.** A wide range of compounds were applied to MDCK cells prior to their water transport turning on at ~24 hours. Concentrations of compounds are indicated in the legends and values are time-normalized to 1 hour before drug addition. A real-time video of the full well plate is shown in Supplementary Video 2.

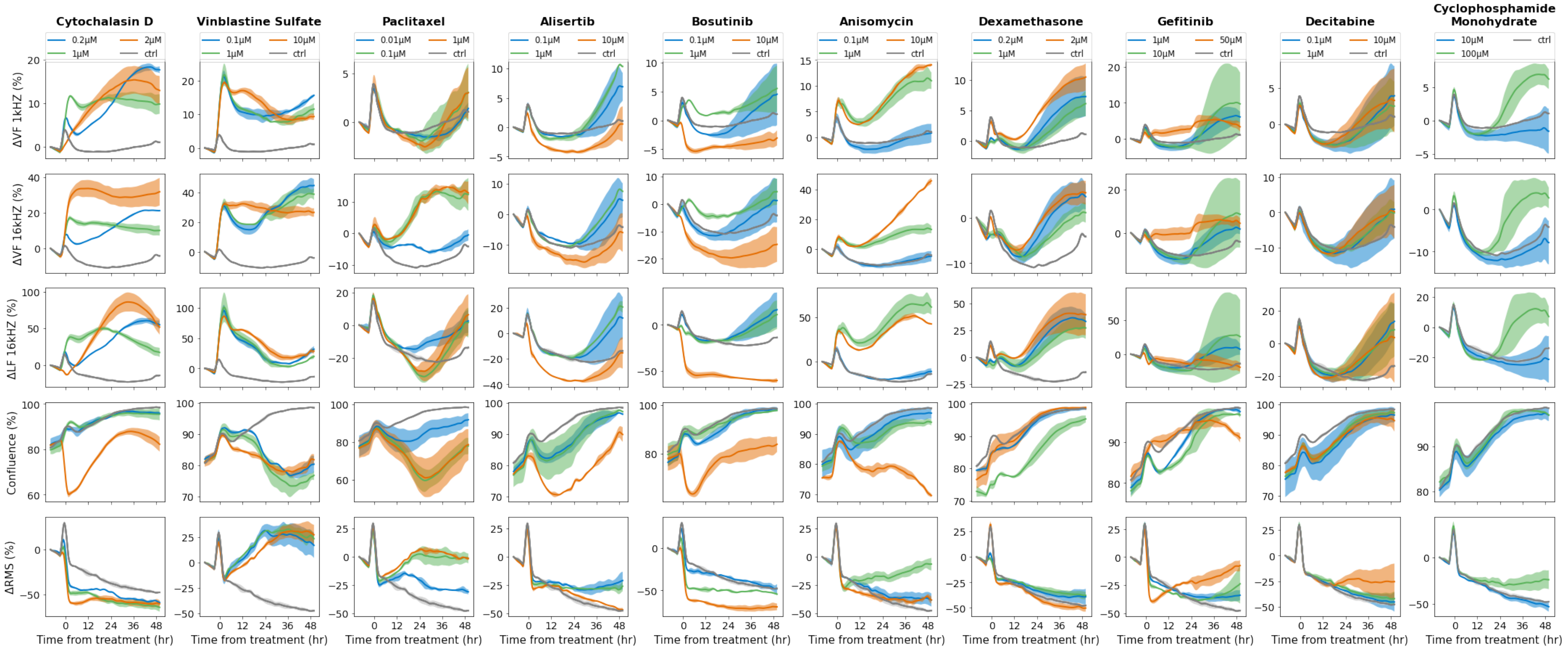


**Supplementary Fig. 9 | Compound applications to A549 cells.** A wide range of drugs were applied to A549 cells ~24 hours after plating. Concentrations of compounds are indicated in the legends and values are time-normalized to 1 hour before drug addition. A real-time video of the full well plate is shown in Supplementary Video 3.

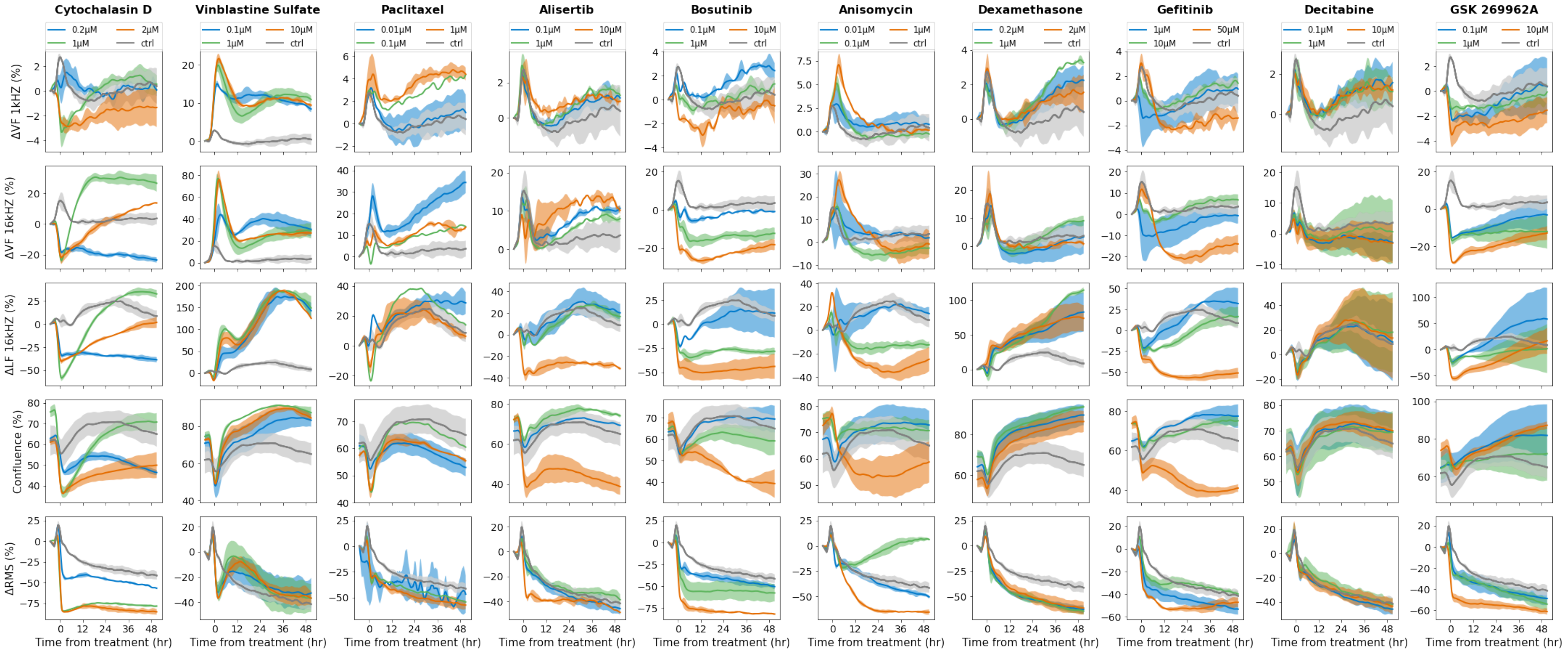


**Supplementary Fig. 10 | Compound applications to MDA-MB-231 cells.** A wide range of drugs were applied to MDA-MB-231 cells ~24 hours after plating. Concentrations of compounds are indicated in the legends and values are time-normalized to 1 hour before drug addition. A real-time video of the full well plate is shown in Supplementary Video 4.

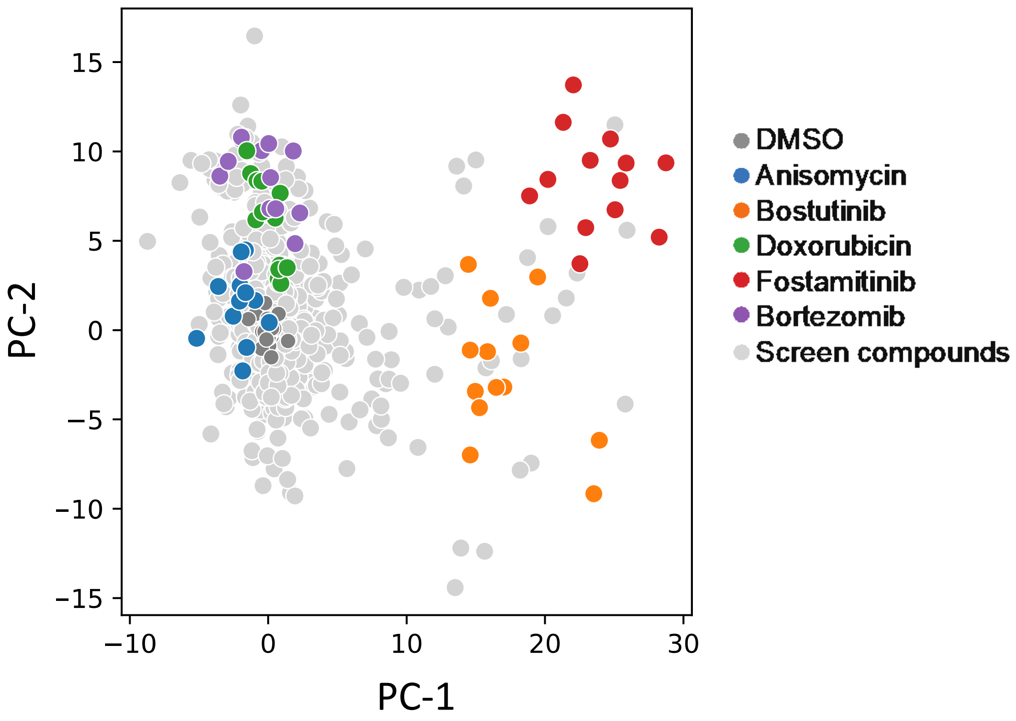


**Supplementary Fig. 11 | Principal component analysis (PCA) of the proof-of-concept diversity screen in Fig. 6.** PCA plot of all points from the 904 compound screen with DMSO (negative control) and 5 positive controls highlighted as indicated in legend.

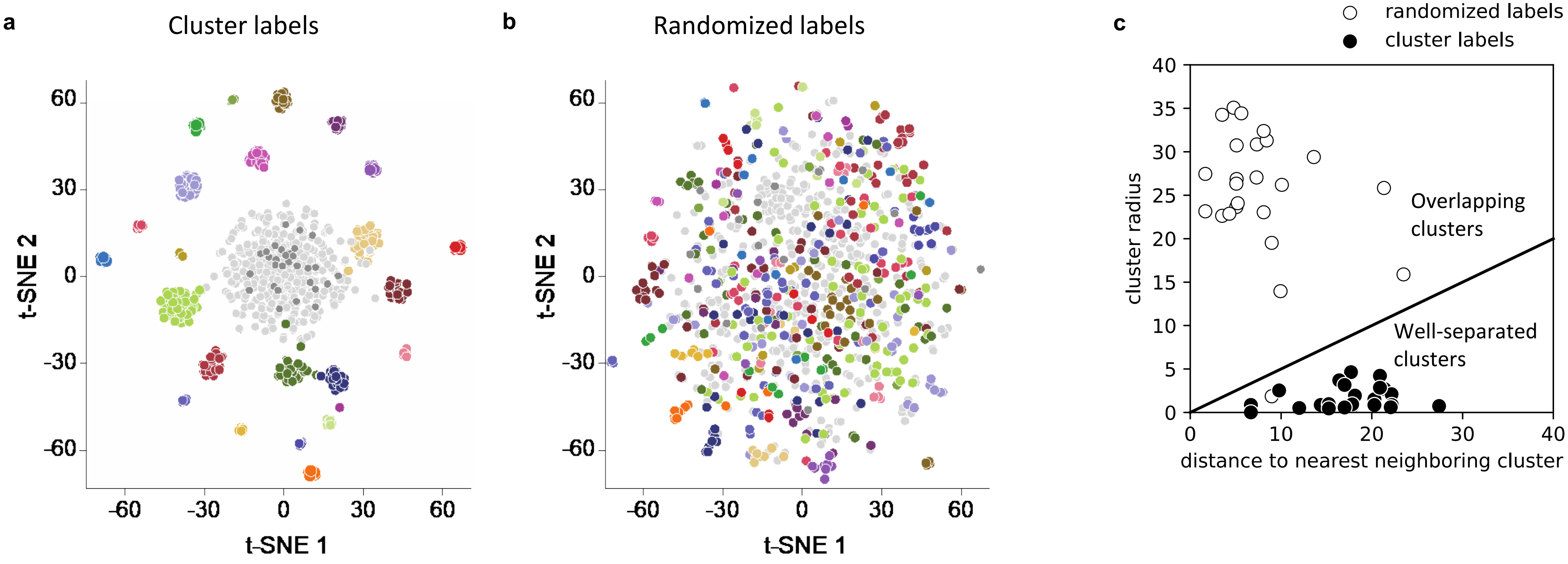


**Supplementary Fig. 12 | Linear discriminate analysis (LDA) model validation.** LDA was performed on the screen dataset of Fig. 6 using labels acquired from clustering algorithm **(a)**, as well as randomizing the cluster labels **(b)**. Using the same LDA parameters, the model was able to clearly distinguish between the datapoints labeled from the clustering algorithm and failed at separating the datapoints with randomized labels. Each color represents a distinct cluster, for 25 total clusters. **c.** Scatter plot of cluster radius vs. distance to nearest neighboring cluster for each individual cluster (25 total). The solid back line denotes the cutoff between overlapping clusters and well-separated clusters (slope of 0.5). Values below 0.5 denote well-separated clusters (as is seen with the cluster-labeled set) with an average slope of 0.11, values above 0.5 denote overlapping clusters (as is seen with the randomized set) with an average slope of 5.04.

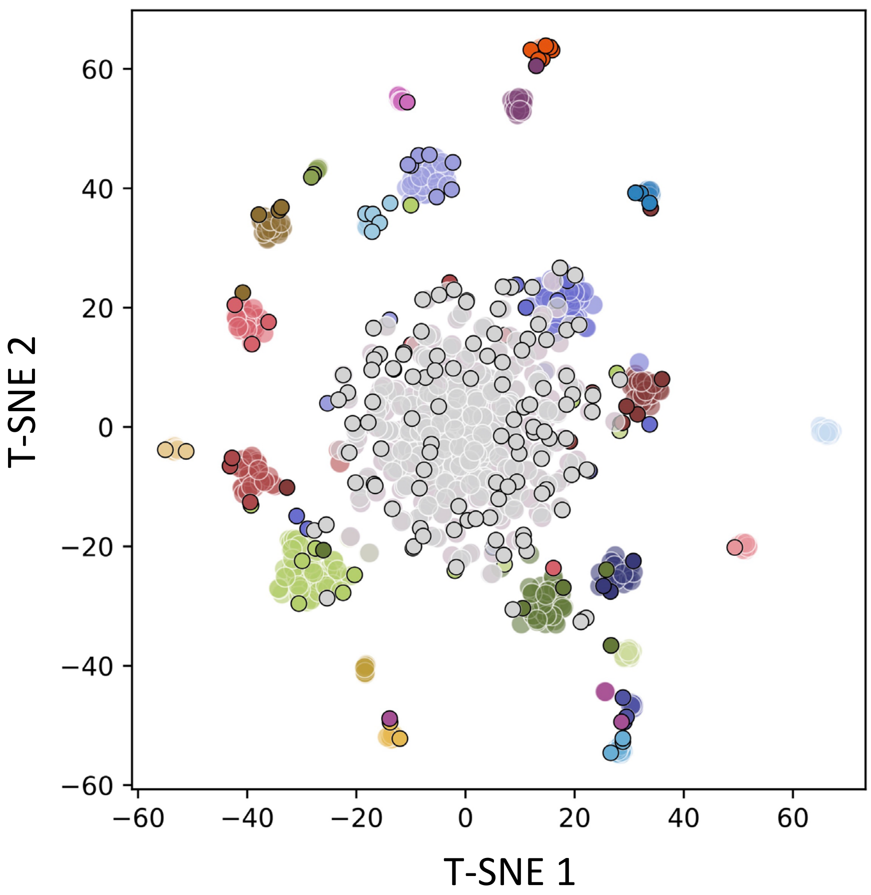


**Supplementary Fig. 13 | Assessment of LDA model performance.** Cluster-labeled 904-compound-screen dataset split into a training (80% of the compounds) and test (20% of the compounds) sets. The model was trained using the training set, predictions were made on the test set with an accuracy score of 0.68

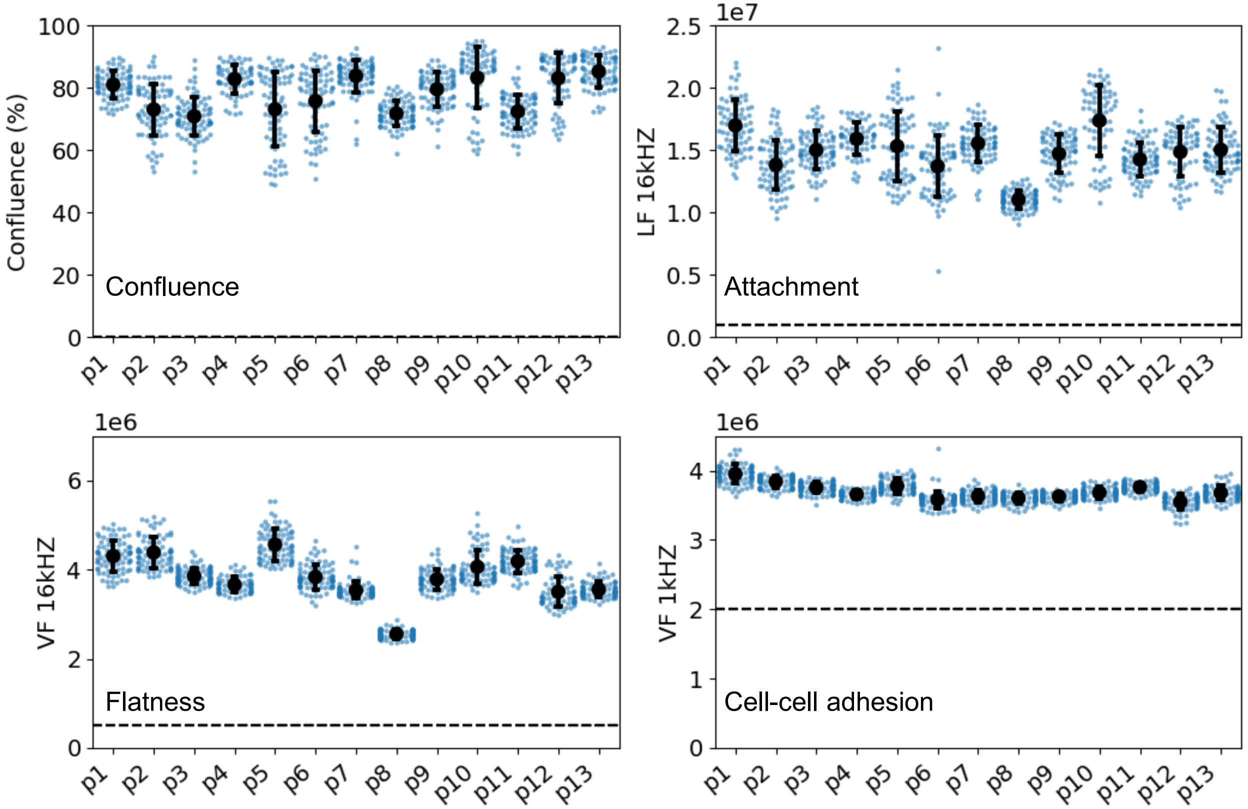


**Supplementary Fig. 14 | Characteristics of A549 cultures before compound additions across technical replicates.** Key parameters were characterized at 24 hours across semiconductor 96-microplates before compound additions. All 13 plates are from the screen presented in Fig. 6 and Supplementary Figs. 11-13. Mean ± s.e for all the wells (points) per plate represented. The dashed line represents the bare electrode impedances before cells were added, showing dynamic range of the field measurements.

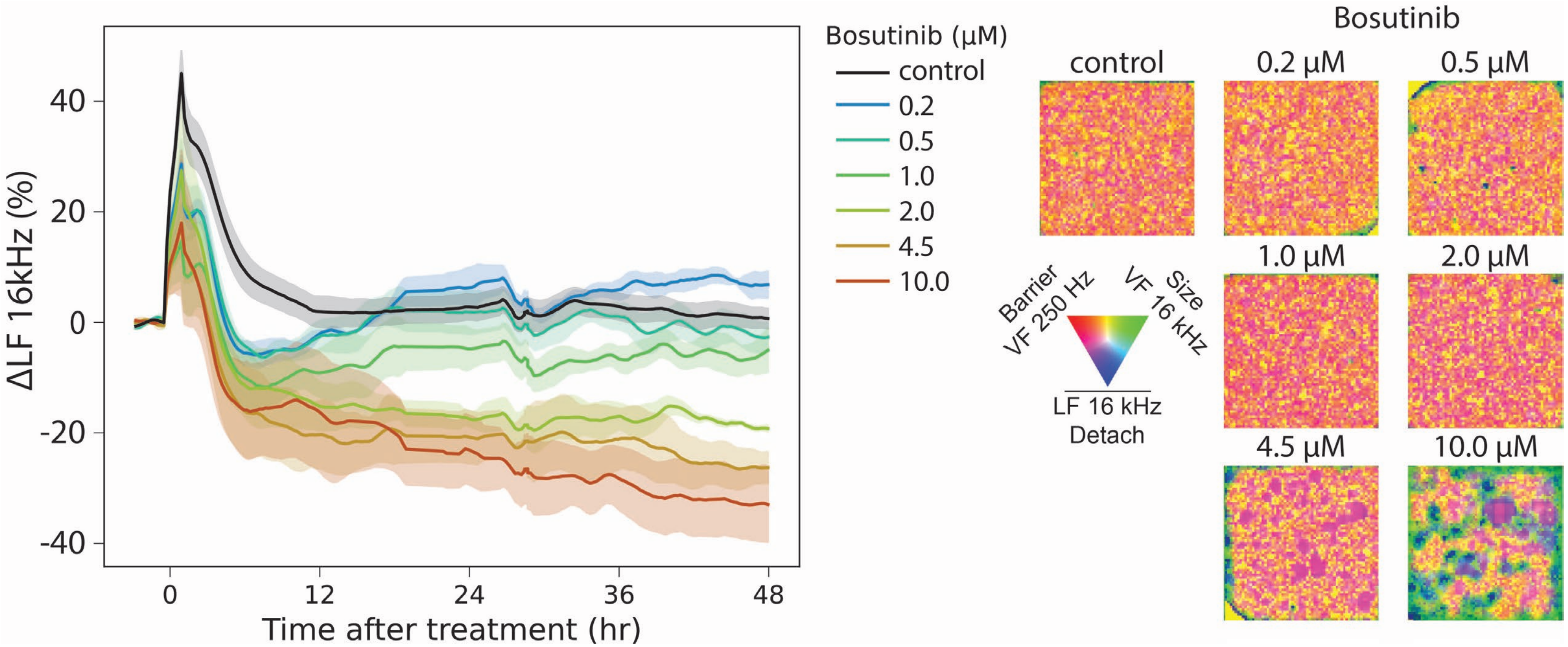


**Supplementary Fig. 15 | Observation of Caco-2 cell doming after Bosutinib treatment on uncoated surface.** A titration of Bosutinib was performed on Caco-2 cells plated on an uncoated surface. A rapid decrease in attachment (left) was observed and domes appeared in impedance images at 40 hours post compound addition (right) for 4.5 μM and 10 μM. Line traces represent mean ± s.d. for 3 wells per condition.

### Supplementary Results

A series of compounds were applied to MDCK, A549 and MDA-MB-231 cells (Supplementary Figs. 8-10). The compounds chosen (Cytochalasin D, Vinblastine Sulfate, Paclitaxel, Alisertib, Bosutinib, Anisomycin, Dexamethasone, Getfitinib, Decitabine, Cyclophosphamide Monohydrate, and GSK 269962A) target various cellular processes including cell division, DNA replication, inflammation and various other signaling pathways. Many diverse changes were observed – increases/decreases in attachment, barrier, cell flatness, motility, and confluency representing cell death/growth – with tight correlation among the three replicates per compound/concentration and dose dependent effects. Many of the effects measured matched known effects of the compounds, and differences in response were observed across cell lines.

As an example of compound modulations observed in these cells, we highlight the effects on motility seen in MDA-MB-231 cells with two different compound treatments: Dexamethasone, an anti-inflammatory drug, and Cytochalasin D, an actin polymerization inhibitor. MDA-MB-231 is established from a metastatic mammary adenocarcinoma and is highly aggressive and invasive[1]. Upon Dexamethasone treatment, an increase in attachment and a decrease in motility (RMS) is observed. Dexamethasone has been described to reverse EMT in MDA-MB-231 cells and reduces their migratory potential[2] corroborating our observed results (Supplementary Fig. 10). A decrease in motility/RMS is also observed upon Cytochalasin D treatment, but in contrast to Dexamethasone, attachment is decreased and there are significant changes to cell flatness – an immediate size decrease for ≥0.2 µM with a subsequent increase for ≥1 µM (Supplementary Fig. 10). Cytochalasin D inhibits actin polymerization, which has been reported to prevent cell motility[2,3] and alter cell shape and flatness[3,4] , supporting our observed results. This further demonstrates that our readout parameters are independent, orthogonal, and measure a wide range of cellular features.

Dexamethasone has also been described to increase barrier in A549 cells. Our measurements show a similar effect of Dexamethasone (Supplementary Fig. 9) that works by increasing the tissue barrier as well as the cell-surface attachment of the cells. Interestingly, in MDCK cells, dexamethasone does not seem to affect any of the morphological parameters (Supplementary Fig. 8). MDCK is a non-cancer cell line with high barrier, low motility, and low levels of inflammation, which might explain the lack of Dexamethasone effect.

In addition to identifying varied effects for compounds, the temporal resolution of the data also enabled us to differentiate between mechanisms of action of drugs with similar outcomes. Paclitaxel and Vinblastine are both microtubule inhibitors, with very similar effects on cell death in A549 cells (Supplementary Fig. 9). At 48 hours post treatment, both compounds show similar levels of cell death, reflected in the confluency measurement. However, in the cell morphology responses between the two drugs reveal distinct effects. The two compounds differ in their mechanism as Paclitaxel is a tubulin polymerization inhibitor, while Vinblastine is a tubulin depolymerization inhibitor. Thus, they have very different effects on cell morphology. The temporal data can thus help differentiate between mechanisms of action.

**Supplementary Video 1: Growth characteristics of eleven cell lines at different densities over 48 hours.** Cells were plated at the densities indicated in Supplementary Figure 6. Densities are laid out from low to high, top to bottom in each column for the indicated cell type. Videos were generated with VF 250 Hz in red, VF 16 kHz in green, and the inverse of the LF 16 kHz in blue. Time ranges from 0-48 hours post seeding.

**Supplementary Video 2: Real-time compound responses in MDCK cells.** MDCK cells were plated and treated with the indicated compounds and concentration in triplicate replicates corresponding to traces shown in Supplementary Figure 8. Cells were seeded at time $t$ = 0, compounds were added at time $t$ = 27 hours. Effects measured up to 48 hours post compound addition. Videos were generated with VF 250 Hz in red, VF 16 kHz in green, and the inverse of the LF 16 kHz in blue.

**Supplementary Video 3: Real-time compound responses in A549 cells.** A549 cells were plated and treated with the indicated compounds and concentration in triplicate corresponding to traces shown in Supplementary Figure 9. Cells were seeded at time $t$ = 0, compounds were added at time $t$ = 27 hours. Effects measured up to 48 hours post compound addition. Videos were generated with VF 250 Hz in red, VF 16 kHz in green, and the inverse of the LF 16 kHz in blue.

**Supplementary Video 4: Real-time compound responses in MDA-MB-231 cells.** MDA-MB-231 cells were plated and treated with the indicated compounds and concentration in triplicate corresponding to traces shown in Supplementary Figure 10. Cells were seeded at time $t$ = 0, compounds were added at time $t$ = 26 hours. Effects measured up to 48 hours post compound addition. Videos were generated with VF 250 Hz in red, VF 16 kHz in green, and the inverse of the LF 16 kHz in blue.
